# Epistasis within the MHC contributes to the genetic architecture of celiac disease

**DOI:** 10.1101/002485

**Authors:** Benjamin Goudey, Gad Abraham, Eder Kikianty, Qiao Wang, Dave Rawlinson, Fan Shi, Izhak Haviv, Linda Stern, Adam Kowalczyk, Michael Inouye

**Affiliations:** NICTA Victoria Research Lab, The University of Melbourne, Parkville, Victoria 3010, Australia; Department of Computing and Information Systems, The University of Melbourne, Parkville, Victoria 3010, Australia; IBM Research, Australia, Level 5, 204 Lygon Street, Carlton, Victoria 3206, Australia; Centre for Systems Genomics, School of Biological Sciences, The University of Melbourne, Parkville, Victoria 3010, Australia; Department of Pathology, The University of Melbourne, Parkville, Victoria 3010, Australia; Department of Mathematics, University of Johannesburg, PO Box 524, Auckland Park 2006, South Africa; Faculty of Medicine, Bar Ilan University, Safed, Israel; Center for Neural Engineering, The University of Melbourne, Parkville, Victoria 3010, Australia

## Abstract

Epistasis has long been thought to contribute to the genetic aetiology of complex diseases, yet few robust epistatic interactions in humans have been detected. We have conducted exhaustive genome-wide scans for pairwise epistasis in five independent celiac disease (CD) case-control studies, using a rapid model-free approach to examine over 500 billion SNP pairs in total. We found 20 significant epistatic signals within the HLA region which achieved stringent replication criteria across multiple studies and were independent of known CD risk HLA haplotypes. The strongest independent CD epistatic signal corresponded to genes in the HLA class III region, in particular *PRRC2A* and *GPANK1/C6orf47,* which are known to contain variants for non-Hodgkin’s lymphoma and early menopause, co-morbidities of celiac disease. Replicable evidence for epistatic variants outside the MHC was not observed. Both within and between European populations, we observed striking consistency of epistatic models and epistatic model distribution. Within the UK population, models of CD based on both epistatic and additive single-SNP effects increased explained CD variance by approximately 1% over those of single SNPs. Models of only epistatic pairs or additive single-SNPs showed similar levels of CD variance explained, indicating the existence of a substantial overlap of additive and epistatic components. Our findings have implications for the determination of genetic architecture and, by extension, the use of human genetics for validation of therapeutic targets.

## Introduction

The limited success of genome-wide association studies (GWAS) to identify common variants that substantially explain the heritability of many complex human diseases and traits has led researchers to explore other potential sources of heritability (in the wide sense), including the low/rare allele frequency spectrum as well as epistatic interactions between genetic variants [1,2]. Many studies are now leveraging high-throughput sequencing with initial findings beginning to elucidate the effects of low frequency alleles [3-6]. However, the characterization of the epistatic component of complex human disease has been limited, despite the availability of a multitude of statistical approaches for epistasis detection [7-13]. Large-scale systematic research into epistatic interactions has been hampered by several computational and statistical challenges mainly stemming from the huge number of variables that need to be considered in the analysis (>100 billion pairs for even a small SNP array), the subsequent stringent statistical corrections necessary to avoid being swamped by large number of false positive results, and the requirement of large sample size in order to achieve adequate statistical power.

The strongest evidence for wide-ranging epistasis has so far come from model organisms [14,15] and recent evidence has demonstrated that epistasis is pervasive across species and is a major factor in constraining amino acid substitutions [16]. Motivated by the hypothesis that epistasis is commonplace in humans as well, recent studies have begun providing evidence for the existence of epistatic interactions in several human diseases, including psoriasis [17], multiple sclerosis [18], Behçet’s disease [19], type 1 diabetes [20], Crohn’s disease [21], bipolar disorder [11] and ankylosing spondylitis [22], as well as complex traits such as serum uric acid levels [23] and the expression levels of multiple genes in human peripheral blood [24]. While these studies have been crucial in demonstrating that epistasis does indeed occur in human disease, several questions remain including how wide-ranging epistatic effects are, how well epistatic pairs replicate in other datasets, how the discovered epistatic effects can be characterized in terms of previously hypothesized models of interaction [25,26], whether it is possible to detect epistatic signal in the presence of strong marginal signals [27,28], and how much (if at all) epistasis contributes to disease heritability [29].

Celiac disease (CD) is a complex human disease characterized by an autoimmune response to dietary gluten. CD has a strong genetic component largely concentrated in the MHC region, due to its dependence on the HLA-DQ2/DQ8 heterodimers encoded by the HLA class II genes *HLA-DQA1* and *HLA-DQB1* [30]. The genetic basis of CD in terms of individual SNP associations has been well characterized in several GWAS [31-34], including the additional albeit smaller contribution of non-HLA variants to disease risk [35]. The success of GWAS for common variants in CD has recently been emphasized by the development of a genomic risk score that could prove relevant in the diagnostic pathway of CD [36]. Autoimmune diseases have so far yielded the most convincing evidence for epistatic associations [37], potentially due to power considerations since these diseases usually tend to depend on common variants of moderate to large effect within the MHC. Given these findings in conjunction with recent observations that rare coding variants may play a negligible role in common autoimmune diseases [3], we sought to determine whether robust epistasis is detectable in CD and whether it accounts for some of the unexplained disease variance.

Here, we present a large-scale exhaustive study of pairwise epistasis in celiac disease. Leveraging Genome-Wide Interaction Search (GWIS), a highly efficient approach for epistasis detection [38], we conduct genome-wide scans for all epistatic pairs across five separate CD case/control datasets of European descent, finding thousands of statistically significant pairs despite stringent multiple testing corrections. Next, we show a high degree of concordance of these interactions across the datasets, demonstrating that they are highly robust and replicable. We characterize the common epistatic models found and compare them to previously proposed theoretical models. Further, given complex linkage disequilibrium patterns, we distil the epistatic pairs down to those that are independent of known HLA risk haplotypes and independent of other epistatic pairs. Finally, we examine whether epistatic pairs add more predictive power and explain more disease variation than additive effects of single SNPs.

## Results

Datasets are summarized in **Table S1**, these include five independent, previously published GWAS datasets of CD with individuals genotyped from four different European ethnicities: United Kingdom (UK1 and UK2), Finland (FIN), The Netherlands (NL) and Italy (IT) [32,33]. To limit the impact of genotyping error and other sources of non-biological variation, we implemented three stages of validation and quality control (QC): (i) standard QC within each dataset, (ii) independent exhaustive epistatic scans within each of the five datasets, and (iii) derivation of a validated list of epistatic interactions based on UK1. The study workflow is shown in **Figure S1**.

### Exhaustive epistatic scans and replication

For each dataset, we implemented stringent sample and SNP level quality control (**Methods**), and then conducted an exhaustive analysis of all possible SNP pairs using the GWIS methodology [38]. Each pair was tested using the GSS statistic, which determines whether a pair of SNPs in combination provides significantly more discrimination of cases and controls than either SNP individually (**Methods**). Forty-five billion pairs were evaluated in the UK1 study (Illumina Hap300/Hap550) and 133 billion SNP pairs were evaluated in each of the four remaining cohorts (Illumina 670Quad and/or 1.2M-DuoCustom). Given this multiple testing burden, we adopted stringent Bonferroni-corrected significance levels of *P* = 1.1 x 10^-12^ for the UK1 and *P* = 3.75 × 10^-13^ for the remaining datasets. Examination of the distribution of observed GSS p-values relative to the uniform p-value distribution showed some deviation (**Figure S2**), indicating the GSS statistic is overly liberal for p-values >10^-5^ yet overly conservative for p-values <10^-5^. We therefore used a permutation-based approach to adjust the observed p-values in a manner analogous to the widely used genomic control method (**Methods**). The resulting adjusted p-value distribution showed no test statistic inflation (**Figure S2**).

To further ensure that the downstream results were robust to technological artefact and population stratification, we took two additional steps: (a) utilizing the raw genotype intensity data available for UK1 for independent cluster plot inspection of 696 SNPs comprising candidate epistatic pairs, and (b) replicating the epistatic interactions of the SNPs passing cluster plot inspection, where replication is defined as a SNP pair passing Bonferroni-adjusted significance both in UK1 and in at least one additional study. Using these criteria, we found that 5,454 SNP pairs (comprising 581 unique SNPs) from the UK1 dataset passed both (a) and (b) above. We denote these pairs as ‘validated epistatic pairs’ (VEPs) below. The full list of VEPs is given in **Dataset S1**. Notably, all VEPs fulfilling these robustness criteria were within the MHC.

More than 134,000 unique pairs achieved Bonferroni-adjusted significance in at least one of the five studies, with the vast majority lying within the extended MHC region of chr 6 (**Figure 1** and **Table S2**). Of the 35 epistatic pairs outside the MHC that were significant in at least one study, none passed Bonferroni-adjusted significance in at least one other study and were thus deemed not replicated. As expected, the number and significance of epistatic interactions increased with sample size. Interestingly, some of the strongest epistatic interactions tended to be in close proximity though few SNPs were in LD with only 1% of pairs having *r*^2^ >0.5 (**Figure S3)**. The heatmaps in **Figure 1** also showed that epistasis was widely distributed with distances of >1Mb common between epistatic pairs. While epistatic interactions were consistently located in and around HLA class II genes, further examination of the VEPs found that many of the strongest epistatic pairs were in HLA class III loci, >1Mb upstream of *HLA-DQA1* and *HLA-DQB1* (**Figure S4**).

**Figure 1:**
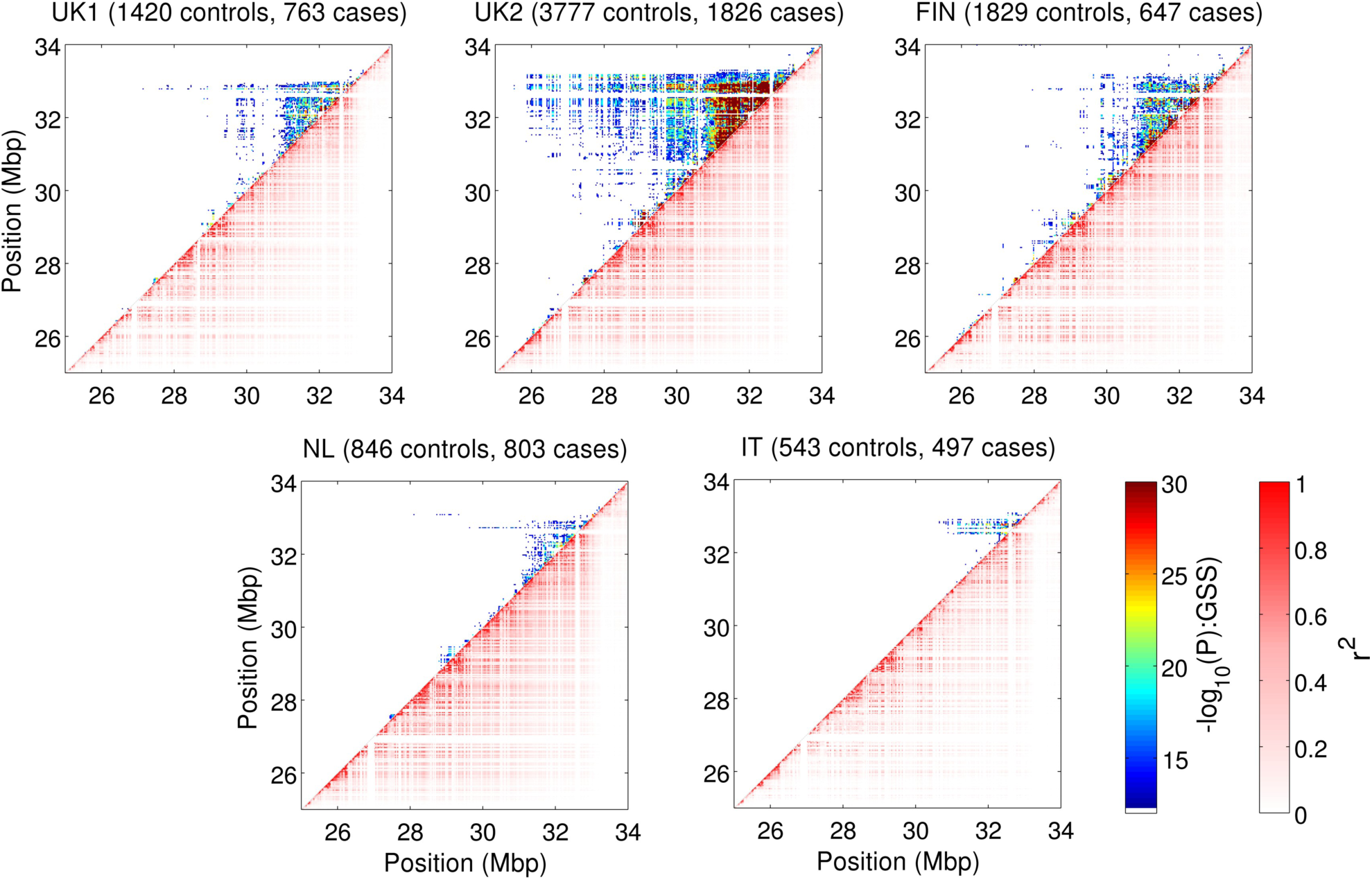
Epistatic interactions and LD patterns within the extended MHC region. SNP pairs within 30KB of each other are shown as a single point on each heatmap. The colour of each point in the upper left half of the graph represents the most significant-log_10_(P-value) returned by the GSS statistic for SNPs pairs within each point. The adjusted-log_10_(P-value) is capped at 30 to increase contrast of lower values. The bottom right half of the graph shows the maximum r^2^ obtained for any SNP pair within a given 30Kb block, demonstrating the strong LD patterns known to exist within this region.

The extent of replication of the epistatic pairs was apparent from the high degree of similarity in the rankings when pairs were sorted by GSS significance (**Figure S5**), with the top 10,000 pairs exhibiting ∼70-80% overlap between the UK1 and UK2 datasets and 40-60% overlap of the UK1 with the pairs found in the NL and FIN datasets. Such high degrees of overlap have essentially zero probability of occurring by chance. The pairs found in the IT dataset showed lower levels of consistency with those detected in the UK1 dataset but overall were still far more than expected by chance with ∼30% overlap at ∼30,000 pairs.

### Independence of Validated Epistatic Pairs and known HLA risk haplotypes

The large number of VEPs demonstrates that thousands of epistatic SNP pairs can reliably be found in the HLA region, however, while the GSS test is designed to select VEPs exhibiting epistatic effects, this does not a priori guarantee that these pairs are not tagging known HLA haplotypes or tagging the same causal signal. Due to the VEPs’ co-localization within a region of complex linkage disequilibrium and the presence of *HLA-DQA1/DQB1* risk haplotypes, known to be instrumental in CD etiology, it is important to estimate the number of epistatic signals that are (a) independent of the known HLA risk haplotypes, and (b) independent of other epistatic signals.

To determine whether VEPs were independent of known haplotypes we utilized a second filtering step in the form of a likelihood ratio test (LRT), testing whether adding a VEP to a logistic regression model of case/control status on the haplotypes increased model fit significantly, with the threshold for significance defined using a false discovery rate (FDR) of 5% (**Methods**). Due to the extensive LD in the region, it is inevitable that some VEPs will be fully captured by HLA risk haplotypes; as a subsequent filter to the GSS test, the LRT will identify these as well as determine whether each VEP contains significantly more independent information on CD risk than the HLA risk haplotype alone. After keeping the VEPs that were independent of risk haplotypes, we used an LD pruning based approach based on Hill’s *Q*-statistic, a normalized chi-squared statistic for multi-allelic loci [39], to estimate the number of haplotype-independent epistatic pairs that were also independent of each other.

From the 5,454 VEPs, we found that 4,744 pairs (>88%) were independent of the known CD risk haplotypes (*DQB1*0302, DQB1*0301, DQB1*0202, DQB1*0201 and DQA1*0301, DQA1*0505, DQA1*0501, DQA1*0201*). From the 4,744 VEPs independent of risk HLA haplotypes, LD pruning using a cutoff of Hill’s *Q* = 0.3 identified 20 VEPs in UK1 which represent independent epistatic signals (**Methods** and **Table S3**).

We further demonstrated the independence of VEPs by employing a LRT evaluating whether the interaction effects for the VEPs holds using a logistic regression-based approach as well as conditioning on known celiac haplotypes and strong univariate associated SNPs (**Methods**). A logistic-regression based test for interaction will differ from the model-free GSS-based test, as it will only detect interactions relative to the log-odds scale while GSS may detect interactions regardless of scale. Nevertheless, we find 1041 VEPs are significant past Bonferroni correction (*P* < 9.12 × 10^-6^) using a meta-analysis based approach **(Methods, Figure S6**) with many showing p-values below 10^-12^. Despite this approach testing for a different definition of interaction and potential loss of power due to overadjustment, many robust epistatic signals remain highly significant.

### Empirical epistatic model distributions

Epistatic models, a subset of two-locus disease models, are typically represented as a table of penetrance values with one penetrance value for each genotype combination [7] and provide insight into how disease risk is distributed. Such models are of interest due to their natural role in the inference of disease mechanism for two or more genes as well as inference of population specific, and therefore potentially evolutionary, effects thereof. Further, characterization of epistatic model frequencies also enables the development of more powerful statistical approaches that target frequent models. While Table S3 shows the models for each of the epistatic pairs chosen to represent the independent signals detected, we are interested in the overall model consistency and distribution across all cohorts and have therefore analysed epistatic models across all VEPs. Following the conventions of Li and Reich [25], we discretized the models for the VEPs to use fully-penetrant values where each genotype combination implies a complete susceptibility or protective effect on disease (**Methods**), simplifying the comparison of models between different SNP pairs.

To establish model consistency, we first replicated the most frequent full penetrance VEP models in the other datasets (**Figure 2**). When considering the distribution of epistatic models we found striking consistency of the UK1 models with those from UK2 and the other Northern European populations (Finnish and Dutch). Only four models from the possible 50 classes [25] occurred with >5% frequency in the Northern European studies, and there was substantial variation in epistatic model as a function of the strength of the interaction. Amongst all VEPs in UK1, the four models corresponded to the threshold model (T; 38.3% frequency), jointly dominant-dominant model (DD; 31.1%), jointly recessive-dominant model (RD; 16.5%), and modifying effect model (Mod; 1.0%) [25,40]. The DD and RD models are considered multiplicative, the Mod model is conditionally dominant (i.e. one variant behaves like a dominant model if the other variant takes a certain genotype), and the T model is recessive. The T model was the most frequent model, especially amongst the strongest pairs.

**Figure 2:**
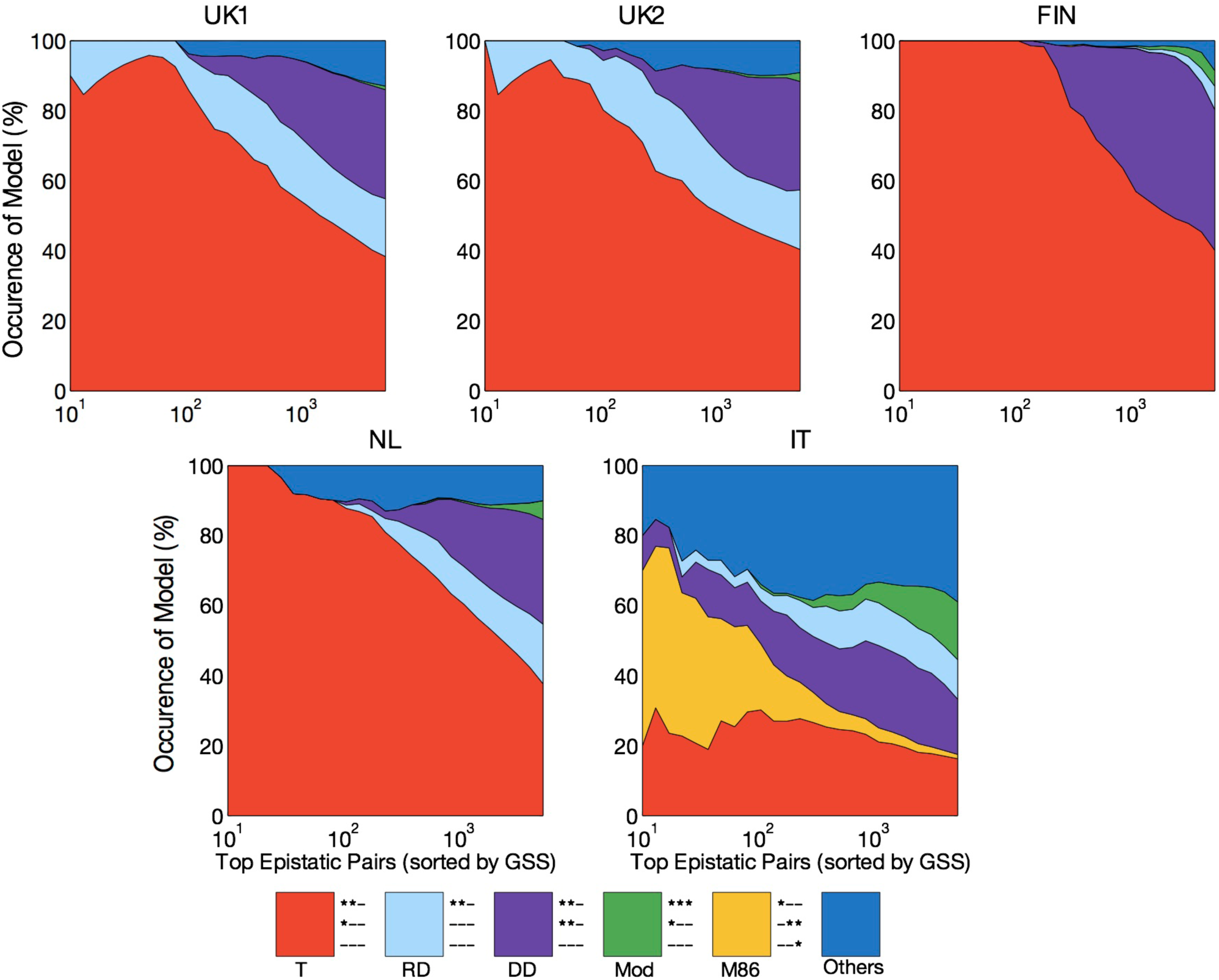
Variation in epistatic models within and between populations. Distribution of epistatic models for VEPs in different studies as increasingly less significant SNP pairs are examined. Different colours represent a different subset of epistatic models. The “other” group represents the remaining set of models. Models have been simplified using the rules provided in [25].

Interestingly, despite the consistency of MHC epistasis, the VEPs showed noticeable differences in epistatic model distribution in the IT dataset. This was in contrast to the other Northern European populations but consistent with the different ranking in GSS significance observed above. In the IT dataset, the distribution of models was altered such that there was a more even distribution. The four most frequent models were still the T model (16.2%), modifying effects (16.5%), DD (15.8%), and RD model (11.3%). But, we also observed that many of the strongest pairs within the IT dataset followed the M86 model, though M86 represented only a small proportion of models overall (1.2%). The other VEP models overall were relatively uniform amongst the remaining models.

The cause(s) of the differences in epistatic model distribution for the IT data are not clear, however it is unlikely due to sample size. While cryptic technical factors cannot be ruled out at this stage, we speculate that there may be population specific epistatic variation that follows the known North/South European genetic gradient [41]. Such variation in epistasis has been previously shown in the evolution of complex genetic systems [42], however evidence of such phenomena in human genetics has not, to our knowledge, been uncovered. While the differences in epistatic model for the IT population relative to Northern Europeans are notable, further studies specifically evaluating the consistency of epistatic model variation between populations (and potentially between diseases) are required to validate its evolutionary basis.

### Contribution of epistatic pairs to celiac disease variance

We next sought to estimate the CD variance explained by the detected epistatic pairs and single SNPs. To do this, we utilized a multivariable model framework which accounted for all SNPs and/or VEPs at once. To assess the contribution of epistatic pairs to CD prediction and thus genetic variance explained, we employed L1 penalized linear support vector machines (SVM, see **Methods**), an approach which models all variables concurrently (single SNPs and/or pairs) and which has been previously shown to be particularly suited for maximizing predictive ability from SNPs in CD and other autoimmune diseases [36,43]. We have previously shown that additive models of single SNPs explain substantially more CD variance than haplotype-based models [36], thus we employ only the former to estimate the gain in CD variance explained here.

We assessed CD variance explained by constructing three separate models: (a) genome-wide single SNPs only, (b) the VEPs only, and (c) a ‘combined’ model of both single SNPs and VEPs together. The models were evaluated in cross-validation on the UK1 dataset, and the best models in terms of Area Under the Curve (AUC) were then taken forward for external validation in the other four datasets without further modification.

In UK1 cross-validation, the combined models led to an increase of ∼1.6% in explained CD variance, from 32.6% to 34.2% (respective, AUC of 0.882 and 0.888) (**Table 1**). In external validation, the models based solely on VEPs had overall high predictive ability across all external validation datasets (AUC > 0.83), but slightly less than models based on single SNPs alone. The combined models yielded the highest externally validated AUC of all models, showing gains in AUC over single SNPs of +0.6% AUC in UK2 (Delong’s 2-sided test *P* = 0.0016). In the IT dataset, average gains were higher at +1.2% AUC yet marginally significant (*P* = 0.0527), and the differences in FIN and NL were smaller (0.5% and 0.1%, respectively) and not significant.

**Table 1:**
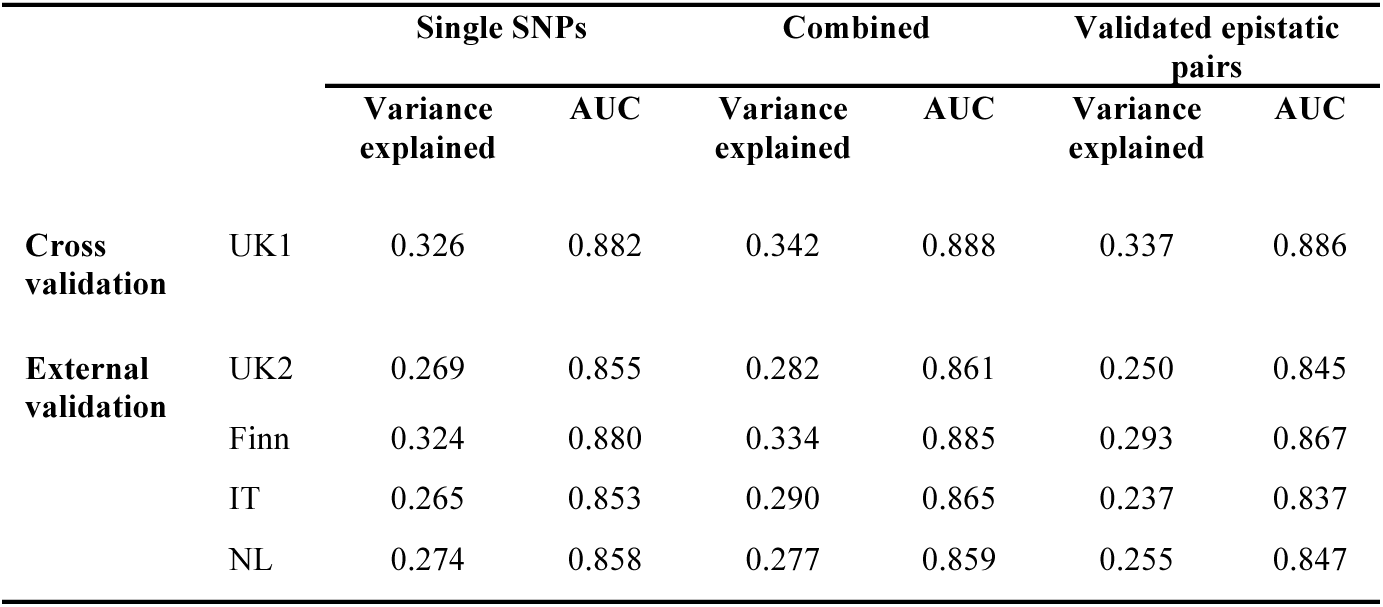
Disease variance explained by models with additive and epistatic genetic effects. Predictive power and disease variance explained by single SNPs and VEPs in cross-validation and in external validation, using SparSNP models. Models were optimized on the UK1 dataset (n=2183 samples) in cross-validation (290K SNPs), and tested without modification on the other datasets. The proportion of disease variance explained (on the liability scale) assumes a population prevalence of 1%. Two-sided DeLong significance tests for AUC of single SNPs+pairs difference from AUC of single SNPs: UK2 *P*=0.001651, FIN *P*=0.2743, IT *P*=0.05271, NL *P*=0.695.

|  |  | Single SNPs |  | Combined |  | Validated epistatic pairs |  |
| --- | --- | --- | --- | --- | --- | --- | --- |
|  |  | Variance explained | AUC | Variance explained | AUC | Variance explained | AUC |
| **Cross validation** | UK1 | 0.326 | 0.882 | 0.342 | 0.888 | 0.337 | 0.886 |
| **External validation** | UK2 | 0.269 | 0.855 | 0.282 | 0.861 | 0.250 | 0.845 |
|  | Finn | 0.324 | 0.880 | 0.334 | 0.885 | 0.293 | 0.867 |
|  | IT | 0.265 | 0.853 | 0.290 | 0.865 | 0.237 | 0.837 |
|  | NL | 0.274 | 0.858 | 0.277 | 0.859 | 0.255 | 0.847 |

Combining the UK1 and UK2 into a single dataset (N=7,786 unrelated individuals) and retraining the models in cross-validation showed similar trends with the best models being the combined models (**Table S4**). The combined models from the larger UK1+UK2 dataset also showed higher AUC in external validation than the models trained on UK1 data only when validating the single SNP + VEP models on the FIN and NL datasets (AUCs +1.3% and +1%; *P* = 0.0007083 and *P* = 0.01113, respectively); however, performance on the IT dataset did not differ significantly (−0.9% AUC, *P*= 0.09568

## Discussion

This study has shown the robust presence of epistasis in celiac disease. The epistatic SNP pairs were mostly independent of HLA risk haplotypes for CD and strongly replicate across cohorts in terms of significance, ranking, and epistatic model. To our knowledge, this level of epistatic signal strength, number of epistatic pairs, and degree of replication has not been previously shown in a complex human disease. We also performed a large-scale empirical characterization of the epistatic models underlying the interactions in CD, with the majority of the VEPs approximately following the threshold model, and a smaller number following dominant-dominant, dominant-recessive, and recessive-recessive models. Further, these patterns were found to be strongly consistent across most of the datasets.

Despite observations that epistatic interactions between SNPs within a locus are enriched for batch effects and poorly clustered genotype clouds [44], the stringent quality control and extensive replication in this study indicate that these SNPs are largely *bona fide* epistatic pairs. A large number of candidate epistatic SNP pairs that did not achieve Bonferroni significance criteria for replication were still highly statistically associated with CD consistently across datasets, indicating that our estimates of the degree of epistasis in CD may be conservative.

For validated epistatic pairs (VEPs), we found that much of the strongest signal was >1Mb upstream of the well-known *HLA-DQA1* and *HLA-DQB1* risk loci and suggested a potentially important epistatic contribution from HLA class III genes. Indeed the strongest epistatic signal, which was independent of HLA risk haplotypes and other VEPs, was attributable to variants in *PRRC2A* and *GPANK1/C6orf47*. Given that individuals with celiac disease are at elevated risk of non-Hodgkin’s lymphoma (NHL), it is intriguing that variants within *PRRC2A* are also associated with NHL [45]. However, for the top *PRRC2A* SNP for NHL (rs3132453) we did not observe a validated epistatic relationship nor linkage disequilibrium between rs3132453 and the epistatic *PRRC2A* SNP (rs2260000), which was low in the HapMap2 CEU (*r*^2^ = 0.05). There is also evidence to suggest that women with celiac disease are at increased risk of early menopause [46,47]. A recent genetic association study of menopausal age identified a missense variant within *PRRC2A* (rs1046089) which was predicted as both structurally damaging for PRRC2A as well as an expression QTL for multiple genes [48]. In our study, the *PRRC2A* SNP rs1046089 showed strong epistasis with another proximal variant in *ABHD16A* as well as several other variants in the MHC despite low LD with the strongest *PRRC2A* epistatic variant (*r*^2^ = 0.27). Overall, our findings indicate that, in addition to the known HLA risk haplotypes for CD, there is epistasis between HLA class III loci, which may have implications for CD co-morbidity.

The epistatic variants were further shown to increase CD variance explained, findings which were replicated in external datasets. Interestingly, models of only epistatic pairs explained nearly as much CD variance as additive models of single SNPs. This observation adds to an increasing body of literature supporting the existence of shared information between additive and epistatic effects [24,49]. In explaining some of the controversy between the contribution of these apparently different classes of effects to a complex disease, our study emphasizes the difficulty in deconvoluting additivity and epistasis, and supports the development of more sophisticated statistical methods to jointly model these effects as well as larger and more powerful studies in which to do so.

While the challenge of deconvoluting additivity and epistasis does not affect studies in the genomic prediction of complex diseases [36], it implies that determining causal genetic signals of CD, and perhaps other autoimmune/inflammatory diseases with substantial HLA-based effects, is more difficult than widely appreciated. Determining the true genetic effects is central to the identification of the molecular products involved in pathogenesis, thus the usage of genetic data in the discovery and validation of therapeutic targets [50] would benefit from the investigation and potential resolution of the epistatic and additive components of a given disease. While the increase in explainable variance from epistatic signals in this study was low, candidate interactions discovered through pairwise analysis may still provide important therapeutic targets [51]. Given this, and in conjunction with recent findings that support the evolutionary persistence of substantial non-additive effects [27], our findings should also stimulate debate around whether the usage of the principle of parsimony (Occam’s Razor) is an adequate rationale for models of exclusively additive effects [29].

These findings have implications for both the genetic architecture of celiac disease as well as the incorporation of epistasis into genetic models of complex disease. The limitations of the first generation GWAS approach to explain missing heritability has led to the development and application of more sophisticated approaches to resolve this problem, yet success has been elusive. Recent results suggest that rare variants add little to known heritability for a number of autoimmune diseases including celiac disease [52]. Epistasis may offer both additional explained (broad-sense) heritability as well as new biology, as evidenced by our findings for HLA class III epistasis. The genetic models of CD generated in this work indicate that while epistatic pairs explain substantial disease variance, overall this variance is largely shared with that of additive effects. Combined epistatic and additive models likely constitute the best solution.

## Methods

### Quality control

A range of quality-control measures were applied to all datasets to limit the impact of genotyping error. For all datasets, we removed non-autosomal SNPs, SNPs with MAF <1%, missingness >1% and those deviating from Hardy-Weinberg Equilibrium in controls with *P* < 5 × 10^-6^. Samples were removed if data missingness was >1%. Cryptic relatedness was also stringently assessed by examining all pairs of samples using identity-by-descent in PLINK, and removing one of the samples if 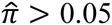. The cryptic relatedness filter removed 17 samples within the UK1 cohort that related to other UK1 samples, and 1208 samples from the UK2 cohort which were either related to other UK2 samples or UK1 samples. Dataset sizes in **Table S1** are reported after the quality control steps above. Significant epistatic SNP pairs were further assessed by manually inspecting the genotyping cluster plots of both SNPs in the UK1 cohort. Intensity data for the other studies was not available. Cluster plot inspection removed 115 SNPs with poor genotyping assays.

### Epistasis detection

The Gain in Sensitivity and Specificity (GSS) test was employed to detect epistasis. The test has been presented in detail in [38] and is summarized in the Supplementary Text. It is available at https://github.com/bwgoudey/gwis-stats.

Analogously to odds ratios used for analyses of single SNPs, we define odds ratios for epistatic pairs based on the GSS statistic

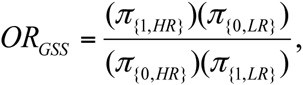

where π_(i,j)_ denotes the proportion of samples with phenotype *i*, 1 for cases and 0 for controls, and carrying genotype combinations which are marked as *j* with *HR* (high risk) indicating genotypes which are associated by GSS with cases and *LR* (low risk) indicating genotypes which are associated with controls (analogous to MDR-style approaches in [12,53]]). By relying on the model-free GSS approach, this odds ratio can be seen as a measure of association for the combination of genotypes from a given SNP pair which has the strongest improvement over the pair’s SNPs.

### Representation of the epistatic models

We approximate the epistatic models for VEPs using two representations: balanced penetrance models and full penetrance models. Following Li and Reich [25] we employ the penetrance, that is, the probability of disease given the genotype, estimated from the data for each of the nine genotype combinations as (number of cases with combination) / (number of individuals with combination). Representing the epistatic model in terms of penetrance allows us to clearly see which genotype combinations contribute more to disease risk (or conversely, may be protective). We employ a standardization to ensure that the penetrance is comparable across datasets, termed *balanced sample penetrance*, and defined as

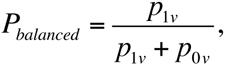

where *p*_*iv*_ refers to the proportional frequency of genotype *v* in class *i*, where controls are 0 and cases are 1. The definition is easily extended to the case of pair of SNPs using the 3 × 3 = 9 possible genotype combinations from each SNP-pair.

For comparison of models, we employ a coarse-grain approach where these values are discretized into binary values, so called “fully penetrant” models, similar to Li and Reich [25]. Unlike Li and Reich, we do not swap the high and low risk status, as we are interested in distinguishing between protective and deleterious combinations. In addition, we do not swap risk status, therefore there will be 100 possible full-penetrance models. For rare genotype combinations we used a simple heuristic, denoting all cells with a frequency below 1% in both cases and controls as ‘low risk’. Experiments with this threshold revealed that altering this cut-off between 0% and 7% made little difference to the overall distribution of our models.

### Independence of epistatic signals from known risk haplotypes

CD strongly depends on specific heterodimers, most notably HLA-DQ2.2, HLA-DQ2.5, and HLA-DQ8, which are in encoded by haplotypes involving the *HLA-DQA1* and *HLA-DQB1* genes, with close to 100% of individuals with CD being positive for one of these molecules. To statistically impute unphased HLA haplotype alleles, we utilized HIBAG [54]. To evaluate whether each VEP was also independent of known CD risk haplotypes, we employed the likelihood ratio test, comparing two logistic regression models: (i) a logistic regression of the phenotype on the risk haplotypes (*DQA1*0201*, *DQA1*0501*, *DQA1*0505*, *DQA1*0301*, *DQB1*0201*, *DQB1*0202*, *DQB1*0301*, and *DQB*0302*) and (ii) a logistic regression including both the haplotypes and the VEP. The haplotypes were encoded as 8 allele dosages [43] and the VEP was encoded as 8 binary indicator variables. We considered an FDR threshold < 0.05, equivalent to *p* < 0.044, as statistically significant, indicating that adding the VEP to the model increased goodness-of-fit over the haplotypes alone (4,744 of the 5,454 tests were FDR-significant).

In addition, we used a logistic regression-based test for bilinear interaction, conditioning on known HLA haplotypes. We compared two logistic regression models: (i) the marginal SNPs and their interaction and (ii) the marginal SNPs alone. As interactions can be induced if two SNPs are in partial linkage with strong univariately-associated SNPs [55], both models were conditioned on the 8 known risk haplotypes and the top 3 associated SNPs found after conditioning on known risk haplotypes (*rs3129763*, *rs2187668* and, *rs3099844*). Again, the haplotypes were encoded using a dosage encoding and VEPs were encoded as 8 binary indicator variables, while the associated individual SNPs were encoded using 2 binary indicator variables. This regression-based test for interaction is different to the model-free GSS statistic as it will only be able to capture a subset of possible interactions due to scale dependencies [56]. We used meta-analysis (Fisher’s method) to determine significance across the five CD cohorts. This is calculated as 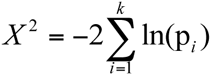 where *k* is the number of independent studies (*k*=5) and *p*_*i*_ is the significance of the LRT each of the cohorts. X^2^ is chi-squared distributed with 2*k* degrees of freedom and hence we can easily derive its significance. Using this meta-analysis approach, 1041 VEPs were significant (after Bonferroni correction *P*= 9.12 × 10^-6^ = 0.05/5454) across the CD cohorts.

### Estimating the number of independent epistatic signals

To determine the number of independent signals coming from the VEPs, we used an LD pruning based approach to filter out all SNP pairs that are in disequilibrium with each other. To ensure that the epistatic signals were not caused by haplotype effects, only SNP pairs that were deemed to be independent of HLA haplotypes were examined. Traditional measures of LD (such as *r*^2^ or *D*’) are designed for examining two binary loci whose frequencies can be reduced to a 2 × 2 table, whereas examining pairs of SNP pairs requires us to examine two multi-allelic loci, whose frequencies are naturally summarized by a 4 × 4 table. A widely used method dealing with this issue is the Hill’s *Q* statistic, a multi-allelic extension of *r*^2^ [39]. It is well known that the *r*^2^ can be derived from the chi-squared test of association, since

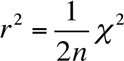

where *n* is the total number of samples and χ^2^ is the chi-squared statistic over the haplotypes formed by the two SNPs. Motivated by this relationship, the *Q* statistic can be expressed as

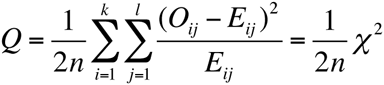

where *O*_*ij*_ and *E*_*ij*_ are the observed and expected haplotype counts when there are *k* and *l* alleles respectively at the two loci. In the case of examining LD between pairs of SNP pairs *k*=*l*=4. Phase information was inferred using SHAPEIT [57], and LD was then computed directly on control samples only. While it is somewhat arbitrary what threshold constitutes independence, given the direct analogy between *r*^2^ and *Q* we utilized a more conservative threshold of *Q* ≤ 0.3 than that commonly used for LD-pruning and tagging procedures (*r*^2^ ≤ 0.5), for example in PLINK. Such conservative thresholds may filter slightly less independent but still informative epistatic signals, thus for the predictive models discussed below, all VEPs were initially allowed to enter the model.

### The predictive models

We have employed a sparse support vector machine (SVM) implemented in SparSNP [58]. This is a multivariable linear model where the degree of sparsity (number of variables being assigned a non-zero weight) is tuned via penalization. The model is induced by minimizing the L1-penalized squared hinge loss

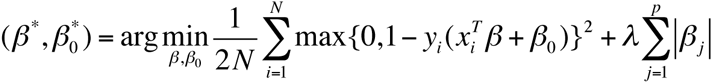

where β and β_0_ are the model weights and the intercept, respectively, *N* is the number of samples, *p* is the number of variables (SNPs and/or encoded pairs), *x*_*i*_ is the *i*th vector of *p* variables (genotypes and/or encoded pairs), *y*_*i*_ is the *i*th case/control status {+1, −1}, and λ ≥ 0 is the L1 penalty. To find the optimal penalty, we used a grid of 100 penalty values within 10 replications of 10-fold cross-validation, and found the model/s that maximized the average area under the receiver-operating characteristic curve (AUC). For models based on single SNPs, we used minor allele dosage {0, 1, 2} encoding of the genotypes. For models based on SNP pairs, the standard dosage model is not applicable; hence, we transformed the variable representing each pair (encoded by integers 1 to 9) to 9 indicator variables, using a consistent encoding scheme across all datasets. The indicator variables were then analyzed in the same way as single SNPs.

### Evaluation of predictive ability and explained disease variance

To maximize the number of SNPs available for analysis, we imputed SNPs in the UK2, FIN, NL, and IT datasets to match those that were in the UK1 dataset but not in former, using IMPUTE v2.3.0 [59]. Post QC this left 290,277 SNPs common to all five datasets. Together with 9×5,454 pairs=49,086 indicator variables, this led to a total of 338,508 markers in the combined singles+pairs dataset. Models trained in cross-validation on the UK1 dataset were then applied without any further tuning to the four other datasets, and the external-validation AUC for these models was then estimated within the validation datasets. To derive the proportion of phenotypic variance explained by the model (on the liability scale), we used the method of Wray et al. [60], assuming a population prevalence of 1%.

## Acknowledgements

MI was supported by a Career Development Fellowship co-funded by the Australian NHMRC and Heart Foundation (#1061435). MI and GA were also supported by University of Melbourne funding. BG, EK, QW, DR, FS, IH and AK were supported by National ICT Australia (NICTA). We thank David van Heel and Cisca Wijmenga for providing the celiac disease data. We thank Karen A. Hunt for cluster plot inspection. We thank Rami Mukhtar for technical advice and Andrew Kowalczyk and Leon Gor for assistance with development of software. We also thank Justin Bedo for helping develop an efficient GSS implementation and Herman Ferra for profiling and processing of statistics. We also thank Armita Zarnegar for assistance with data processing and John Markham, Justin Bedo and Geoff Macintyre for insightful discussions and comments.

